# Mitotic Spindle Positioning (MISP) is an actin bundler that selectively stabilizes the rootlets of epithelial microvilli

**DOI:** 10.1101/2021.08.02.454661

**Authors:** E. Angelo Morales, Cayetana Arnaiz, Evan S. Krystofiak, Marija Zanic, Matthew J. Tyska

## Abstract

Microvilli are conserved actin-based surface protrusions that have been repurposed throughout evolution to fulfill diverse cell functions. In the case of transporting epithelia, microvilli are supported by a core of actin filaments bundled in parallel by villin, fimbrin, and espin. Remarkably, microvilli biogenesis persists in mice lacking all three of these factors, suggesting the existence of unknown bundlers. We identified Mitotic Spindle Positioning (MISP) as an actin binding factor that localizes specifically to the rootlet end of the microvillus. MISP promotes rootlet elongation in cells, and purified MISP exhibits potent filament bundling activity *in vitro*. MISP-bundled filaments also recruit fimbrin, which further elongates and stabilizes bundles. MISP confinement to the rootlet is enforced by ezrin, which prevents decoration of the membrane-wrapped distal end of the core bundle. These discoveries reveal how epithelial cells optimize apical membrane surface area and offer insight on the remarkable robustness of microvilli biogenesis.

## INTRODUCTION

Surface protrusions are essential features that enable cells to interact with the external environment in all domains of life. Choanoflagellates, the closest living relatives of animals, are unicellular eukaryotes that developed the earliest known polarized feeding apparatus consisting of long-lived actin-based protrusions, which we now generally refer to as microvilli (Sebé-Pedrós et al., 2013). Multicellular eukaryotes eventually maximized solute transport by compartmentalizing large numbers of microvilli on the surface of specialized hollow organs (Peña et al., 2016). In animals, striking examples of such organization are found on the apical luminal surface of enterocytes in the small intestine, where densely packed arrays of thousands of microvilli extend from the surface of individual cells, collectively forming the ‘brush border’ (Crawley et al., 2014). In other specialized cases, arrays of microvilli have been repurposed to serve diverse functions including sperm recognition in oocytes, mechanosensation in inner ear hair cells, and light perception in photoreceptor cells, among others (Lange, 2011).

An individual microvillus extends from the cell surface as a finger-like membrane protrusion, supported by a core of 20-40 actin filaments bundled in parallel (Mooseker and Tilney, 1975; Ohta et al., 2012). Core actin bundles exhibit lengths on the micron scale and flexural rigidities high enough to deform the enveloping plasma membrane (Atilgan et al., 2006). Previous biochemical studies established that at least three different bundling proteins assemble the microvillar core bundle: villin, fimbrin (also known as plastin-1) and espin (Bartles et al., 1998; Bretscher and Weber, 1979, 1980). Villin is the first bundler recruited apically during brush border differentiation, followed by fimbrin, and espin (Bartles et al., 1998; Ezzell et al., 1989). Single villin or espin knockout (KO) mice exhibit near-normal microvillar morphology and organization (Ferrary et al., 1999; Pinson et al., 1998; Revenu et al., 2012). In contrast, fimbrin KO mice exhibit microvilli that are ~15% shorter (Grimm-Günter et al., 2009; Revenu et al., 2012). Remarkably, villin-espin-fimbrin triple KO mice are viable and their enterocytes still assemble apical brush borders, although microvillar length is reduced by ~40% (Revenu et al., 2012). This latter finding underscores the remarkable robustness of microvillar growth and further suggests that brush border assembly is driven by multiple factors operating in parallel, some of which remain unidentified.

Within the microvillus, actin filaments that comprise the core bundle are oriented with their barbed ends toward the distal tips and pointed ends extending down into the subapical cytoplasm (Mooseker and Tilney, 1975). The barbed ends are the preferred site of new actin monomer incorporation whereas the pointed ends are the favored site of disassembly (Pollard and Mooseker, 1981). This kinetic difference at opposite ends creates a system that allows subunits to flux retrograde through the core bundle in a process referred to as ‘treadmilling’ (Kirschner, 1980). Indeed, recent studies with epithelial cell culture models revealed that treadmilling is crucial for microvilli assembly and motility (Gaeta et al., 2021; Meenderink et al., 2019).

While most of the core actin bundle protrudes from the cell surface enveloped in plasma membrane, a much shorter segment – the ‘rootlet’ – extends down into the subapical cytoplasm. Core bundle rootlets are directly linked to a dense filamentous network called the ‘terminal web’, an organelle-free zone that presumably regulates trafficking to and from the apical plasma membrane (Mooseker et al., 1983). Ultrastructural studies first suggested that rootlets are interconnected with terminal web filaments at least in part by non-muscle myosin-2 and spectrin (Hirokawa et al., 1982). Deep in the terminal web, rootlets appear to be directly crosslinked with a meshwork of cytokeratins (Hirokawa et al., 1982, 1983). One possible crosslinking factor is the actin bundler fimbrin, which is found along the length of the core actin bundle with an apparent enrichment on the rootlet (Grimm-Günter et al., 2009). Based on the highly interconnected nature of filaments throughout the terminal web, this network likely serves as a physical platform that offers long-term stability and mechanical support for protruding brush border microvilli. Although core bundle rootlets can only interact with the terminal web filaments if they remain free of membrane wrapping, factors that protect the proximal end of the bundle from membrane encapsulation during microvillar growth remain undefined.

Biophysical investigations have also established that the structural stability of microvilli is promoted by tethering core actin bundles to the surrounding plasma membrane (Nambiar et al., 2010). Core bundles are laterally bridged to their enveloping membrane by myosin-1a and −6, as well as ezrin (Berryman et al., 1993; Hegan et al., 2012; Howe and Mooseker, 1983). Recent studies on the dynamics of growing microvilli revealed that ezrin accumulation in a nascent microvillus occurs in parallel with core bundle elongation, and that loss of ezrin from the protrusion leads to microvillus collapse (Gaeta et al., 2021). As one of the most highly characterized membrane-cytoskeleton linkers, ezrin adopts two conformational states: an open ‘active’ state when phosphorylated and a closed ‘inactive’ state when dephosphorylated (Bretscher et al., 1997). Dynamic cycling between these two states allows ezrin to bridge treadmilling actin bundles to the enveloping plasma membrane (Viswanatha et al., 2012), while providing continuous mechanical support for the protrusion. However, mechanisms that constrain ezrin enrichment to the distal segment of the core bundle and control the extent of membrane wrapping remain poorly understood.

Here we report Mitotic Spindle Positioning (MISP) as a novel brush border component that targets specifically to the rootlets of microvillar core actin bundles. Previous studies revealed that MISP is an actin binding protein implicated in promoting mitotic spindle orientation and mitotic progression (Kumeta et al., 2014; Maier et al., 2013; Zhu et al., 2013), although a role in native tissues has yet to be reported. In intestinal epithelial cells, we found that MISP is enriched in the subapical region beneath the plasma membrane at the base of the brush border, where it colocalizes with fimbrin along core bundle rootlets. Loss- and gain-of-function studies revealed that MISP functions to elongate rootlets and limit the extent of membrane wrapping of core actin bundles. Consistent with these phenotypes, MISP bundles actin filaments *in vitro* and in cells, creating structures that are primed for fimbrin recruitment. Overexpression of both factors leads to a striking overgrowth of hyper-stable rootlets from the subapical domain. Further, we found that MISP confinement to microvillar rootlets depends on the presence of active ezrin in the microvillus. Overall, our findings lead to a new model for rootlet specification whereby ezrin and MISP exert mutual exclusivity to establish membrane-wrapped vs. unwrapped segments of the core bundle. MISP confinement to rootlets, in turn, recruits fimbrin to further facilitate crosslinking of the proximal ends of core bundles in the terminal web. This work holds important implications for understanding the assembly and stabilization of actin-based protrusions in diverse epithelial systems, and also provides a molecular rationale for the remarkable robustness of brush border assembly alluded to in previous multigene loss-of-function mouse models (Delacour et al., 2016).

## RESULTS

### MISP localizes to the rootlets of brush border microvilli

In a previous proteomic study, we identified peptides from MISP in brush borders isolated from mouse small intestine (McConnell et al., 2011). To validate MISP as a *bona fide* brush border resident and to examine its localization in native tissues, we immunostained longitudinal paraffin sections of mouse small intestine. Confocal microscopy of stained sections revealed that MISP specifically localizes to the brush border along the full length of the crypt-villus axis (**Fig. 1A**). To label core actin bundles, we stained tissue sections with antibodies directed against villin. Using a membrane marker to delineate the apical surface, we found that MISP is highly enriched in the terminal web and exhibits mutually exclusive labeling with the membrane-wrapped protruding microvilli (**Fig. 1A, B**). As previously reported (Dudouet et al., 1987; Robine et al., 1985), we found that villin signal gradually increased along the crypt-villus axis, following the direction of enterocyte migration and differentiation **(Fig. 1A, C, D; magenta labels)**. In contrast, MISP signal remains relatively constant along the crypt-villus axis **(Fig. 1A, C, D; green labels)**, suggesting that this factor is apically targeted independent of differentiation state. We were able to recapitulate this observation in cell culture using CACO-2BBE cells, which differentiate and take on an enterocyte-like phenotype after a prolonged period of confluent culture (Peterson and Mooseker, 1993; Peterson et al., 1993). In this system, MISP was also expressed and localized at early and late stages of microvilli assembly **(Fig. 1E-G)**. These results indicate MISP is a component of the brush border that is highly enriched in the subapical terminal web throughout enterocyte differentiation.

**Figure 1.**
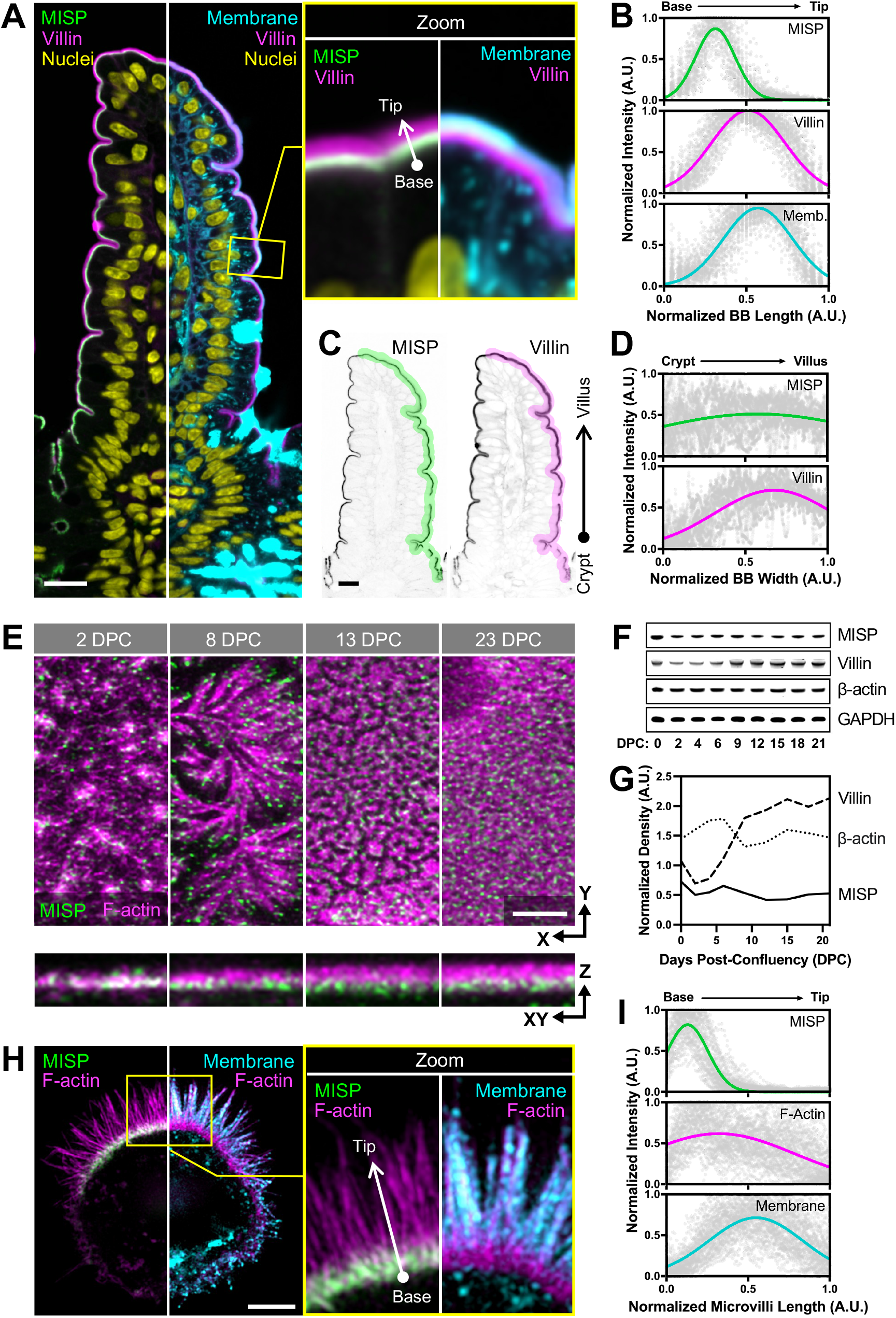
MISP localizes to the rootlets of brush border microvilli. **(A)** Confocal images of a small intestinal tissue sections stained for MISP (green), villin (magenta), DNA with DRAQ5 (yellow), and membrane with WGA (cyan). Main panel shows a split image to facilitate visualization of distinct three-color merge combinations. The upper right panel shows zoomed images of the region in the main panel highlighted by the yellow box. Scale bar = 15 *μ*m. **(B)** Fluorescence intensity distributions of MISP, villin, and a plasma membrane marker measured parallel to the base-tip axis of the brush border (BB), oriented as indicated by the white arrow in the zoomed panel A; n = 200 line scans measured on five small intestinal villi. **(C)** Inverted single-channel images of MISP and villin signals from panel A. Scale bar = 15 *μ*m. **(D)** Fluorescence intensity distribution of MISP and villin measured relative to the crypt-villus axis of the brush border (BB) as represented by the highlighted region in panel C. Black arrow in panel C shows the orientation of scans; n = 11 scans of six small intestinal villi. **(E)** Confocal maximum intensity projection image of CACO-2_BBE_ cells at different stages of differentiation stained for MISP (green) and F-actin with phalloidin (magenta). Upper panels show XY *en face* views of the apical surface at the indicated days post-confluency (DPC) time point, whereas bottom panels show resliced XZ views to facilitate the visualization of MISP localization along the microvillar axis. Scale bar = 3 *μ*m. **(F)** Western blot time series of CACO-2_BBE_ cell lysates probed for MISP, villin, β-actin, and GAPDH at the indicated DPC. **(G)** Density measurements of MISP, villin, and β-actin western blot signals from panel F normalized to GAPDH and plotted as a function of DPC. **(H)** SIM maximum intensity projection image of a W4 cell stained for MISP (green), F-actin with phalloidin (magenta), and membrane with WGA (cyan). Left panel shows a split image to facilitate visualization of distinct two-color merge combinations; zoom panel shows the region highlighted by the yellow box. Scale bar = 3 *μ*m. **(I)** Fluorescence intensity distributions of MISP, Factin, and apical membrane from line scans measured parallel to the microvillar axis, oriented as indicated by the white arrow in the zoomed panel H; n = 58 line scans of single core actin bundles from 10 cells. All plots in panels B, D and I show Gaussian curve fits of the raw data points for each channel.

Given that MISP is a highly specific terminal web component with previously described actin binding potential (Kumeta et al., 2014), we sought to determine if MISP associates with rootlets at the base of microvilli. To examine this possibility in more detail, we turned to LS174T-W4 cells (herein referred to as ‘W4 cells’), a human intestinal epithelial cell line that has been engineered to provide switch-like control over brush border assembly (Baas et al., 2004). Using super-resolution structural illumination microscopy (SIM), we found that MISP was highly enriched on the rootlet segments of core bundles that extend immediately beneath the apical membrane (**Fig. 1H, I**), which was consistent with the localization we observed in mouse intestinal tissue. We also examined the localization of an EGFP-tagged variant of MISP expressed in LLC-PK1-CL4 cells, a pig kidney epithelial cell culture model that allows for visualization of individual microvilli extending from the apical surface (Gaeta et al., 2021). In these cultures, we again noted a striking enrichment of MISP on microvillar rootlets, with signal that was mutually exclusive with the membrane-wrapped protruding segment of the core bundle (**Fig. S1A, B**). Together, these localization studies in native tissues and multiple cell culture models uniformly indicate that MISP specifically targets to core bundle rootlets and is excluded from the membrane-wrapped segment of the microvillus.

### MISP is required for maintaining rootlets at the base of microvilli

Core bundle rootlets are anchored in the terminal web, which likely provides mechanical support for brush border assembly and longterm stability. To determine if MISP is required for normal brush border assembly and microvillar structure, we generated W4 cell lines with stable shRNA-mediated knockdown (KD) of MISP. Loss of MISP was confirmed by Western Blot analysis **(Fig S2A, B)**. Using low magnification confocal microscopy, we scored the fraction of cells that were brush border positive as indicated by polarized F-actin staining. At a population level, we found that the percentage of W4 cells forming a polarized brush border dropped from 82% in the scramble control to 70% in MISP KD cells **(Fig. 2A, B)**. This modest phenotype was rescued when an EGFP-MISP construct refractory to KD was reintroduced **(Fig. 2B)**. However, the overall intensity of F-actin per cell decreased significantly in MISP KD cells (**Fig. 2C**), suggestive of a marked perturbation in Factin network architecture even in cells that still exhibited polarized brush border assembly. To further understand the impact of MISP loss-of-function, we took a closer look at individual MISP KD cells that still formed a polarized brush border. Measurements of microvillar dimensions revealed that the overall length of core actin bundles did not change significantly between scramble control and MISP KD cells (**Fig. S2C**). However, using a membrane marker to delineate the membrane-wrapped vs. unwrapped segments of the core bundle, we found that the protruding microvillus increased in length (1.96 ± 0.37 μm in controls vs. 2.22 ± 0.39 μm in KD) at the expense of rootlet length, which decreased significantly (0.43 ± 0.09 μm in controls vs. 0.27 ± 0.07 μm in KD; **Fig. 2D, E**). Consistently, the membrane coverage of total core actin bundles also increased slightly (82% in controls vs. 89% in MISP KD cells; **Fig. S2D**). Thus, rootlet shortening in MISP KD cells is driven by membrane overwrapping of core bundles.

**Figure 2.**
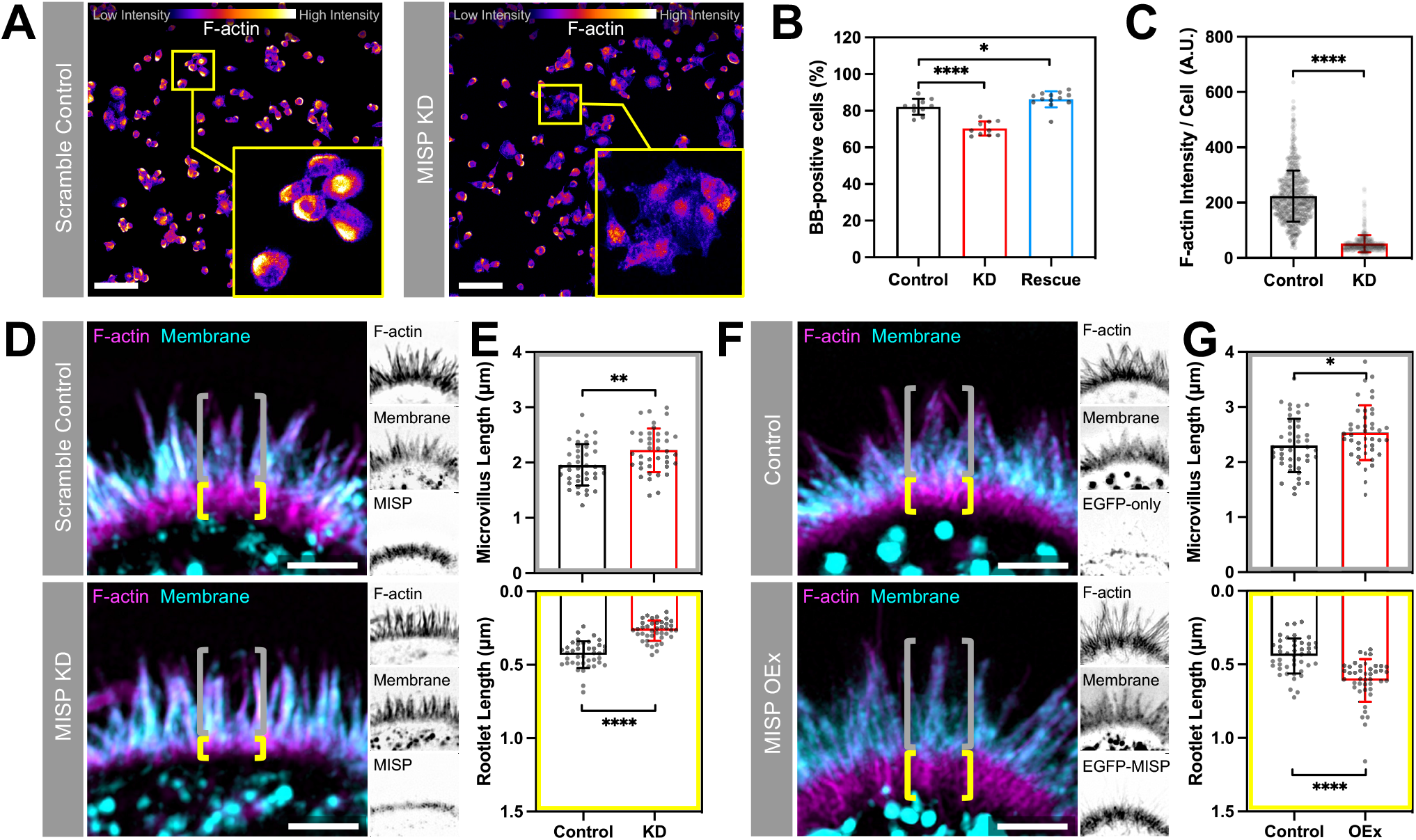
MISP is required for maintaining rootlets. **(A)** Confocal maximum intensity projection images of scramble control and MISP knockdown (KD) W4 cells stained for F-actin; intensity color-coded with ‘Fire’ LUT, where warmer colors denote higher intensities. Zoomed images show representative cells for each condition. Scale bar = 100 *μ*m. **(B)** Percentage of W4 cells forming brush borders (BB-positive cells) from panel A comparing scramble control, MISP KD, and EGFP-MISP rescue conditions. Each data point represents the percentage of BB-positive cells in a single large field of view 620 μm^2^); n ≥ 10 fields per condition. **(C)** F-actin intensity measurements of W4 cells from panel A comparing scramble control and MISP KD conditions. Each data point represents the averaged F-actin intensity of a single cell; n > 600 cells per condition. **(D)** SIM maximum intensity projection images of the brush border of W4 cells in scramble control (top) and MISP KD (bottom) conditions stained for F-actin (magenta) and membrane (cyan); each panel shows merged images to the left and inverted single channel images to the right. Yellow brackets indicate the extension of actin rootlets, whereas gray brackets show extension of the membrane-wrapped segment of the core bundle in both conditions. Scale bars = 2 *μ*m. **(E)** Length measurements of the protruding membrane-wrapped segment of core bundles (‘Microvillus Length’; gray, top plot) and core bundle rootlets (‘Rootlet Length’; yellow, bottom plot) from scramble control and KD cells. Each data point represents the average of > 10 length measurements per cell; n ≥ 40 cells per condition. All data points are representative of three independent experiments. **(F)** SIM maximum intensity projection images of the brush border of W4 cells in control (top) and MISP overexpressing (OEx, bottom) cells stained for F-actin (magenta) and membrane (cyan); each panel shows merged images to the left and inverted single channel images to the right. Yellow brackets indicate the extension of actin rootlets, whereas gray brackets show extent of microvillar protrusion in both conditions. **(G)** Length measurements of the protruding membrane-wrapped segment of microvillus (gray, top) and core bundle rootlets (yellow, bottom) from control and MISP OEx cells. Each data point represents the average of >10 length measurements per cell; n ≥ 40 cells per condition. All data points are representative of three independent experiments. All bar plots and error bars denote mean ± SD. p-values were calculated using the unpaired T-test (*: p < 0.05; **: p < 0.01; ****: p < 0.0001).

As loss of MISP shortened rootlets and increased membrane wrapping on core bundles, we sought to determine if increasing MISP levels would elongate rootlets at the expense of membrane wrapping. To test this hypothesis, we overexpressed EGFP-MISP in W4 cells and examined microvillar structure using super-resolution microscopy. Similar to the localization studies described to above, EGFP-MISP exhibited specific enrichment on microvillar rootlets. Relative to control cells, MISP overexpression promoted a significant elongation of both the membrane-wrapped (2.30 ± 0.48 μm in controls vs. 2.54 ± 0.49 μm in OEx) and rootlet (0.44 ± 0.12 μm in controls vs. 0.61 ± 0.15 μm in OEx) segments of the core bundle (**Fig. 2F, G)**. Consistently, this resulted in a significant increase in the overall length of core actin bundles, and a slight reduction in membrane coverage of total core actin bundles in MISP-overexpressing cells **(Fig. S2E, F)**. Taken together, these findings show that MISP promotes microvillar rootlet elongation and protects the proximal end of the core bundle from membrane wrapping.

### Purified MISP assembles tightly packed linear actin bundles *in vitro*

Based on the terminal web localization of MISP and the impact of MISP perturbation on rootlet length, we sought to determine if purified MISP is sufficient to drive the formation of linear F-actin bundles similar in structure to core bundle rootlets. Full length human MISP was reported to be highly insoluble in previous purification attempts (Kumeta et al., 2014), and thus far only truncated fragments have been studied *in vitro*. To aid with solubility, we tagged the N-terminus of MISP and EGFP-MISP with a maltose binding protein (MBP) and purified these variants from Sf9 insect cells for further characterization (**Fig. S3A, B**). Using a low-speed sedimentation assay, we found that MBP-tagged full length MISP robustly bound to and bundled actin filaments in a concentration dependent manner **(Fig. 3A)**, with a bundling affinity (K_D_) of 0.23 mM (**Fig. 3B**). To directly visualize the impact of MISP on F-actin organization and bundling, we mixed MBP-EGFP-MISP with phalloidin-stabilized actin filaments and then examined the resulting structures using confocal microscopy. MISP/F-actin mixtures exhibited extensive bundling and crosslinking of filaments, particularly in regions that were heavily decorated with MBP-EGFP-MISP **(Fig. 3C**; **zoom 1)**. F-actin intensity in bundles was also significantly higher when MISP was present in solution **(Fig. 3D)**. Interestingly, we noticed that in some instances, MISP accumulated at the ends of actin bundles where the phalloidin signal was lower (**Fig. 3C, E; red arrowheads in zoom 2)**. To examine the ultrastructural organization of these samples, we turned to transmission electron microscopy (TEM). To reduce the possibility of functional interference from the MBP moiety, we removed this tag using TEV protease before TEM imaging (**Fig. S3C**). In control samples (F-actin alone), TEM images revealed single actin filaments that extended for many microns across the grid surface **(Fig. 3F; left panel)**. In contrast, MISP/F-actin mixtures exhibited extensive crosslinking of filaments and formation of tightly packed linear actin bundles **(Fig. 3F; right panels)**. Although the tightly packed and 3D nature of these bundles precluded clear determination of filament polarity in our images, spacing measurements revealed that filaments in these bundles were separated by an average distance of 10.2 ± 2.5 nm **(Fig. 3G)**, which is shorter than the distances between filaments bundled by villin or espin (~12 nm), but comparable to the spacing produced by fimbrin (Bartles et al., 1998; Hampton et al., 2008; Matsudaira et al., 1983; Volkmann et al., 2001). Together, these findings demonstrate that MISP is sufficient to form tightly packed linear actin bundles with an inter-filament spacing similar to that of fimbrin.

**Figure 3.**
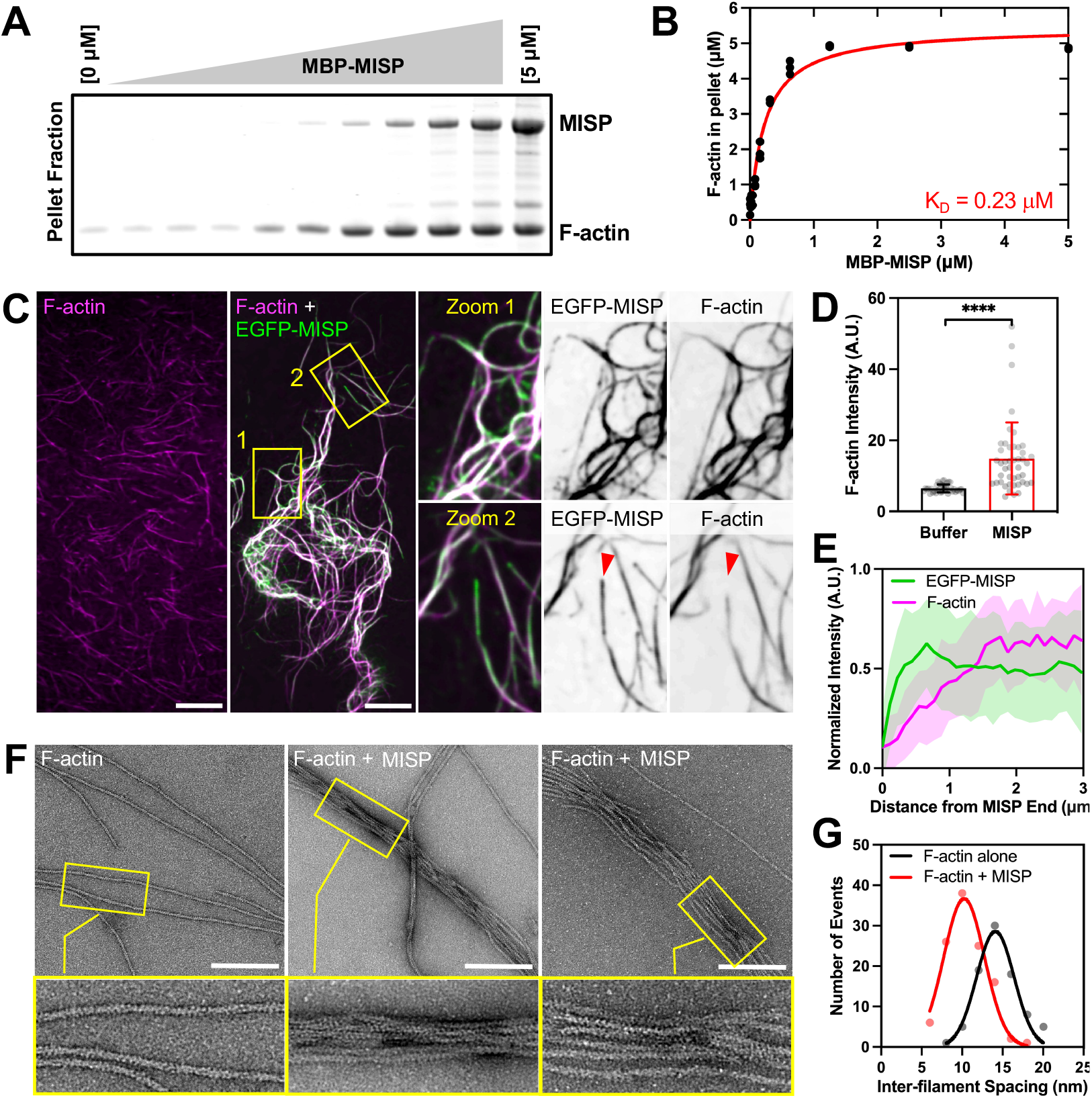
Purified MISP assembles tightly packed linear actin bundles *in vitro*. **(A)** Low-speed sedimentation assay of F-actin stabilized with phalloidin (5 *μ*M) and increasing concentrations of MBP-MISP (0 – 5 *μ*M). Coomassie-stained SDS polyacrylamide gel shows the pellet fractions recovered after centrifugation. **(B)** Density quantification of Coomassie-stained bands shown in panel A. All data points correspond to three independent experiments and were fit using a hyperbolic saturation binding model yielding a K_D_ = 0.23 mM. **(C)** Confocal images of F-actin stabilized with phalloidin (0.5 *μ*M; magenta) alone or pre-mixed with MBP-EGFP-MISP (0.1 *μ*M; green). Zoomed images correspond to the yellow boxes shown in the merged channel; single channels are shown as inverted images. Red arrowheads indicate the end of MISP-bundled actin filaments. Scale bar = 10 *μ*m. **(D)** Fluorescence intensities of F-actin in buffer alone or with MISP from panel C. Each data point corresponds to the integrated intensity value of a 250 μm^2^ field of view; n ≥ 39. Bar plots and error bars denote mean ± SD. p-values were calculated using the unpaired T-test (****: p < 0.0001). **(E)** Line scan analysis of EGFP-MISP (green) and F-actin (magenta) intensities measured at the ends of bundles shown in panel C. **(F)** Transmission electron microscopy images of negatively stained phalloidin stabilized F-actin (0.2 *μ*M) in buffer alone or pre-mixed with purified MISP (0.04 *μ*M). Scale bar = 200 nm. **(G)** Histogram of interfilament spacing measurements from bundles shown in the yellow boxes in panel F. Each data point corresponds to the average of ≥ 110 values; bin size = 2. Averaged values were fit using a Gaussian curve.

### MISP recruits fimbrin to actin bundles

Among the three previously characterized actin bundlers in the brush border, fimbrin is the only one that appears to preferentially accumulate on core bundle rootlets, where it might mediate physical interactions with the terminal web cytokeratin network (Grimm-Günter et al., 2009). We therefore sought to determine if MISP binding to actin depends on fimbrin, either cooperatively or competitively. To this end, we turned to HeLa cells, which do not typically form microvilli but can assemble a variety of other actin-based networks. Interestingly, mCherry-MISP expression alone promoted the formation of aberrant actin bundles throughout the cytoplasm (**Fig. 4A; left panel)**, whereas EGFP-fimbrin expression had little impact on existing actin networks (**Fig. 4A; middle panel)**. However, when MISP and fimbrin were co-expressed, fimbrin was robustly recruited to MISP-bundled actin filaments (**Fig. 4A; right panel)**, where it demonstrated strong colocalization with MISP (**Fig. 4B, C**). These data indicate that MISP promotes the formation of actin bundles, which in turn recruit fimbrin, and further suggest hierarchical functioning of these factors during microvillar assembly.

**Figure 4.**
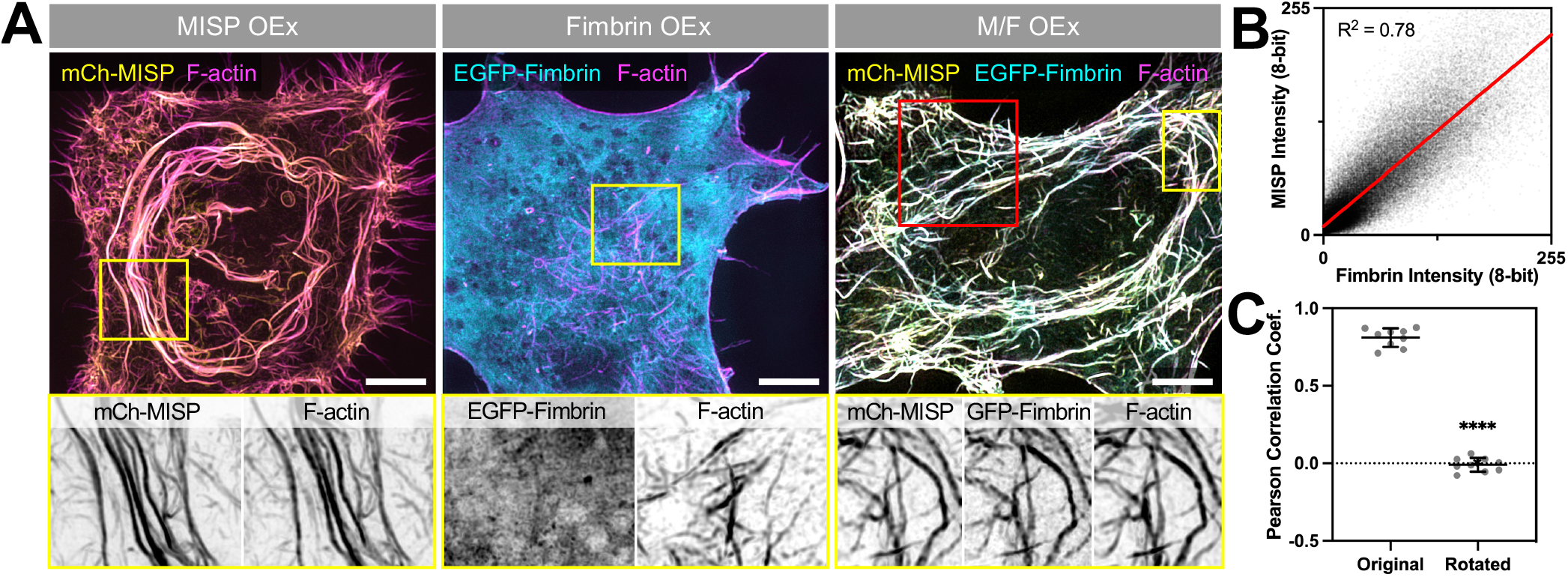
MISP recruits fimbrin to actin bundles. **(A)** SIM maximum intensity projection images of HeLa cells overexpressing mCherry-MISP (left panel), EGFP-fimbrin (middle panel), and mCherry-MISP and EGFP-fimbrin (right panel). All cells were stained for F-actin with phalloidin (magenta). Each panel shows merged channels on top; inverted single channel images along the bottom show zoomed regions highlighted by the yellow box in the merge images. Scale bar = 5 *μ*m. **(B)** Colocalization analysis of mCherry-MISP and EGFP-fimbrin intensities along actin bundles shown in the red box in panel A; data were fit using linear regression. **(C)** Pearson correlation coefficient measurements between mCherry-MISP and EGFP-fimbrin intensities along actin bundles. ‘Original’ denotes aligned raw channels, whereas ‘Rotated’ denotes MISP channel merged with a fimbrin channel that was rotated 90°; loss of correlation following rotation indicates channel overlap is non-random. Each data point represents correlation measurements of a single 15 *μ*m^2^ ROI per cell; n = 9 cells. Error bars denote mean ± SD. p-values were calculated using the unpaired T-test (****: p < 0.0001).

### MISP and fimbrin cooperate to elongate microvillar rootlets

We next sought to determine if MISP and fimbrin cooperate to elongate rootlets in polarized W4 epithelial cells. HALO-MISP and EGFP-fimbrin co-expression resulted in a dramatic hyperelongation of rootlets, which extended deep into the cell (**Fig. 5A**). The tangled nature of these exaggerated rootlets prevented us from measuring the length of individual core bundles in these structures. Instead, we focused on measuring the length of protruding microvilli as well as the maximum distance that rootlets reached into the cytoplasm using a membrane marker as a point of reference. While the length of microvilli increased moderately, the reach of rootlets increased significantly by ~3-fold in cells coexpressing MISP and fimbrin compared to cells overexpressing either MISP or fimbrin alone (3.01 ± 1.35 μm vs. 0.91 ± 0.41 μm vs. 1.18 ± 0.49 μm, respectively) (**Fig. 5A, B**). We also observed that these exaggerated rootlet networks converged as they grew further from the apical membrane (**Fig. 5A, C**). Colocalization analysis showed a strong correlation between MISP and fimbrin signals throughout these structures (**Fig. 5D**). To further define the properties of the exaggerated rootlets promoted by MISP and fimbrin co-expression, we conducted Fluorescence Recovery After Photobleaching (FRAP) assays on W4 cells expressing HALO-β-actin alone, or in combination with EGFP-fimbrin and mCherry-MISP. Photobleaching of the HALO-β-actin signal allowed us to directly interrogate actin dynamics in distinct regions of interest (ROIs) in transfected cells. In the microvilli of control cells, β-actin turned over with a t_half_ of 126.7 s, which likely reflects the treadmilling rate of core bundles in this system (**Fig. 5E, G; green labels**). However, in cells co-expressing MISP and fimbrin, we noted two distinct recovery rates: β-actin in protruding microvilli turned over at a rate that was 4-fold slower than controls (t_half_ = 529.1 s), whereas recovery in exaggerated rootlets was extremely slow to nonexistent (**Fig. 5F, G; magenta and cyan labels, respectively**). Therefore, consistent with their actin bundling activities, MISP and fimbrin co-expression hyper-stabilized and reduced β-actin flux through both rootlets and core bundles in protruding microvilli.

**Figure 5.**
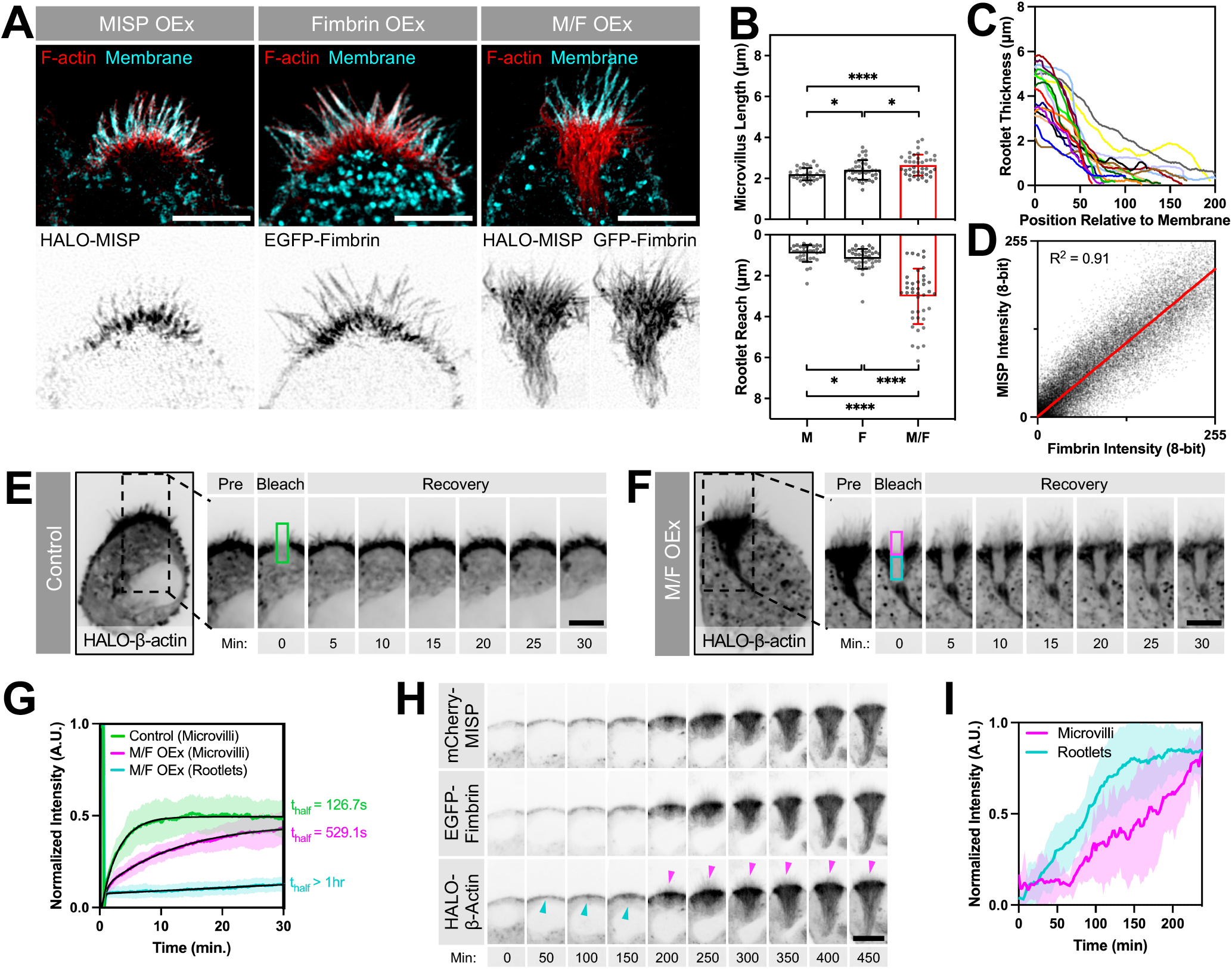
MISP and fimbrin cooperate to elongate microvillar rootlets. **(A)** SIM maximum intensity projection images of W4 cells overexpressing HALO-MISP (left panel), EGFP-fimbrin (middle panel), or HALO-MISP and EGFP-fimbrin together (right panel). All cells were stained for F-actin with phalloidin (red) and membrane with WGA (cyan). Each panel shows merged channels on top with inverted single channel images corresponding to overexpressing constructs along the bottom. Scale bar = 5 *μ*m. **(B)** Length measurements of membrane-wrapped segment of the core bundle (‘Microvillus Length’, top plot) vs. distance that rootlets extend into the cytoplasm (‘Rootlet Reach’, bottom plot) in W4 cells overexpressing the constructs described in panel A. Each data point represents the average length of > 10 length measurements per cell; n ≥ 34 cells per condition. All data points are representative of three independent experiments. Bar plots and error bars are mean ± SD. p-values were calculated using the unpaired T-test (*: p < 0.05; ****: p < 0.0001). **(C)** Rootlet width measurements of W4 cells overexpressing HALO-MISP and EGFP-fimbrin. Width was plotted starting at the membrane boundary (x = 0) extending down into the cell body where rootlet ends converged. Each line represents a measurement from a single cell; n = 10 cells. **(D)** Colocalization analysis between HALO-MISP and EGFP-fimbrin intensities measured along rootlets shown in panel A. Values were fit using linear regression. **(E, F)** Photobleaching analysis of W4 cells overexpressing HALO-β-actin alone (E) or HALO-β-actin with EGFP-fimbrin and mCherry-MISP (F). Although a single ROI was positioned on the brush border and bleached in both conditions, the analysis region in panel F was subdivided into two sub-ROIs to quantify differences in the recovery of the apical microvilli (magenta box) vs. subapical rootlets (cyan box). Scale bar = 5 *μ*m. **(G)** Fluorescence intensity recovery of HALO-β-actin from conditions described in panels E and F; measurements were taken from the color-coded ROIs from panels E and F; n > 14 cells per condition. All intensity values for each condition are shown as mean ± SD. Averaged values for each condition were fit using two-phase association curves. **(H)** Confocal maximum intensity projection time series image montages of W4 cells overexpressing HALO-β-actin, EGFP-fimbrin and mCherry-MISP after adding doxycycline to induce brush border assembly. Cyan arrowheads denote initiation of terminal web actin network assembly. Magenta arrowheads denote microvilli assembly. Scale bar = 10 *μ*m. **(I)** Fluorescence intensity of HALO-β-actin signal during brush border assembly from panel H. ‘Rootlet’ signal was measured from the subapical area; ‘microvilli’ signal was measure from the apical area; n = 10 cells.

To further understand how these hyper-elongated and stable rootlets assemble relative to protruding microvilli, we used live imaging to visualize brush border assembly in W4 cells expressing mCherry-MISP, EGFP-fimbrin and HALO-β-actin (**Movie S1**). During the first two hours after the addition of doxycycline to promote brush border assembly, we first observed the assembly of a terminal web actin network immediately beneath the apical cap **(Fig. 5H, I; cyan labels)**. As this dense network accumulated subapically, microvilli eventually began to emerge **(Fig. 5H, I; magenta labels)**. Consistent with our FRAP analysis, core bundle rootlets elongated from the subapical region below microvillar protrusions with little apparent actin turnover or disassembly from the basal ends **(Fig. 5H, I; β-actin channel)**. These results suggest that the assembly of a terminal web actin network precedes the assembly of microvilli, which is consistent with the proposed role of the terminal web in offering mechanical support for protrusion formation (Tilney and Cardell, 1970).

### MISP and ezrin exhibit mutually exclusive targeting along core actin bundles

Our localization studies in native tissues and cell culture models establish that MISP localizes specifically to the rootlet segment of the core actin bundle, which remains free of plasma membrane wrapping. Although such specific targeting has not been described before, one possible explanation is that MISP is normally prevented from occupying the membrane-wrapped segment of the core bundle by other actin binding factors found at this end of the protrusion. One potential competing factor is ezrin, which functions as a membrane-actin linker that is needed for the structural stability of microvilli (Casaletto et al., 2011; Saotome et al., 2004). Ezrin was also previously reported to regulate MISP levels at the cell cortex in dividing HeLa cells (Kschonsak and Hoffmann, 2018). In W4 cells, we found that endogenous ezrin localizes to the membrane-wrapped ends of core actin bundles as expected, generating a distribution that is mutually exclusive with MISP rootlet labeling (**Fig. S6**).

To determine if MISP confinement to microvillar rootlets is enforced by ezrin, we expressed an EGFP-ezrin construct in W4 cells and monitored the impact on endogenous localization and levels of MISP. In control W4 cells, SIM images revealed that MISP signal was uniformly distributed at the base of the brush border as expected (**Fig. 6A, B; left panels**). However, in W4 cells expressing EGFP-ezrin, we noted that MISP signal was displaced towards the brush border periphery; MISP was almost entirely excluded from the center of the apical domain where EGFP-ezrin levels were highest (**Fig. 6A, B; right panels**). 3D rendering of ezrin-overexpressing cells revealed that MISP signal appeared as a striking ring-like structure surrounding ezrin signal at the center of the brush border (**Movie S2**). When we stained ezrin-overexpressing cells with phalloidin and WGA to visualize F-actin and the plasma membrane, we observed a reduction in F-actin signal in regions lacking MISP signal **(Fig. 6A, B)**. This was also accompanied by drastic shortening of core bundle rootlets (0.43 ± 0.10 μm in controls vs. 0.28 ± 0.06 μm in OEx), and a significant increase in length of protruding microvilli (2.16 ± 0.36 μm in controls vs. 2.58 ± 0.40 μm in OEx; **Fig. 6A, C)**. Interestingly, length measurements of membrane-wrapped vs. unwrapped segments of core bundles in ezrin-overexpressing cells are similar to what we observed in MISP KD cells **(Fig. 2)**. Together these findings indicate that normal levels of ezrin are required to restrict MISP targeting to core bundle rootlets, which in turn promotes their elongation.

**Figure 6.**
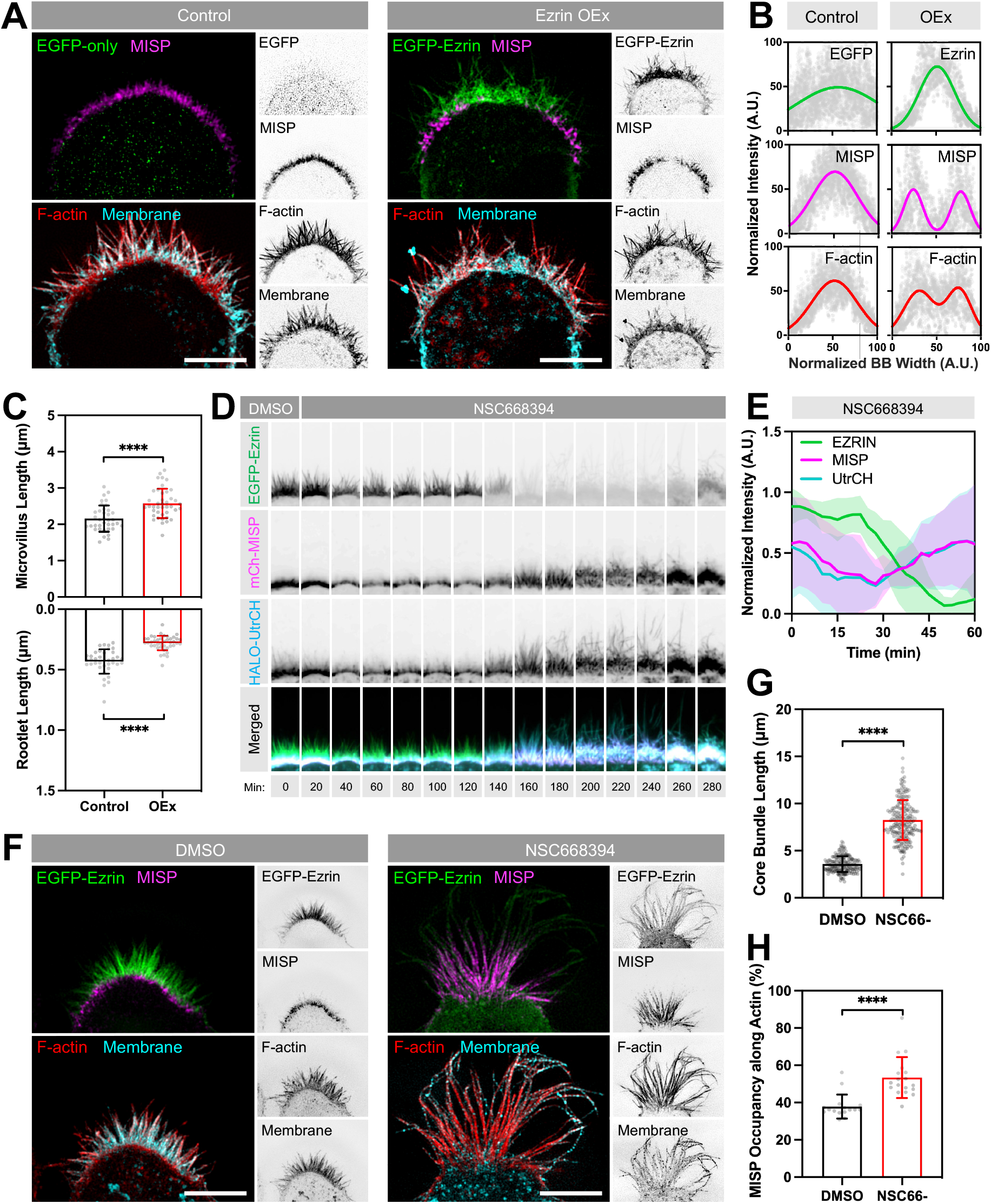
MISP and ezrin exhibit mutually exclusive targeting at opposite ends of core actin bundles. **(A)** SIM maximum intensity projection images of W4 cells overexpressing EGFP alone (green, left panel) or EGFP-ezrin (green, right panel), and stained for endogenous MISP (magenta), F-actin with phalloidin (red), and membrane with WGA (cyan). Each two-color merge image is shown with the corresponding inverted single channel images to the right. Scale bar = 5 *μ*m. **(B)** Intensity distributions across the brush border (BB) from left to right, measured for each of the markers described in panel A. Distributions were fit using single or double Gaussian curves. Number of cells per condition ≥ 8. **(C)** Length measurements of the membrane-wrapped segments of core actin bundles (‘Microvillus Length’, top plot) and core bundle rootlets (‘Rootlet Length’, bottom plot) from W4 cells under the conditions shown in panel A. Each data point represents the average length of > 10 length measurements per cell; n ≥ 37. All data points are representative of three independent experiments per condition. **(D)** Confocal maximum-intensity projection time series image montages of a W4 cell overexpressing EGFP-ezrin (green), mCherry-MISP (magenta), HALO-UtrCH (blue) before and after adding the ezrin inhibitor (NSC668394). The width of each box in the montage is 7 *μ*m. **(E)** Fluorescence intensity measurements of markers described in panel D. Data is shown as mean ± SD. **(F)** SIM maximum intensity projection images of W4 cells overexpressing EGFP-ezrin in DMSO (left panel) or NSC668394 (right panel) conditions. Cells were also stained for endogenous MISP (magenta), F-actin with phalloidin (red), and membrane with WGA (cyan). Each panel shows two-color merge with their corresponding inverted single channel images to the right. Scale bar = 5 *μ*m. **(G)** Length measurements of core actin bundles corresponding to the conditions described in F. Each data point represents the length of a single core actin bundle; n > 190 length measurements. **(H)** Percentages of MISP coverage along core bundles corresponding to the conditions described in panel F. Each data point represents the percentage of microvillar membrane coverage per cell; n > 20 cells. All bar plots and error bars denote mean ± SD. p-values were calculated using the unpaired T-test (****: p < 0.0001).

Within microvillar protrusions, phosphorylated ezrin adopts an open conformational state that bridges the membrane to the actin cytoskeleton (Bretscher et al., 1997). We hypothesized that the open and active conformational state of ezrin within membrane protrusions is responsible for restricting MISP to core bundle rootlets. To test this idea, we used a small molecule inhibitor of ezrin (NSC668394), which disrupts its phosphorylation and actin-binding capacity (Bulut et al., 2012). We overexpressed EGFP-ezrin, mCherry-MISP and HALO-UtrCH (an F-actin binding probe based on the calponin homology domain of utrophin) in W4 cells and monitored their fluorescence intensity over time before and after the addition of NSC668394 (50 μM). Using confocal microscopy, we observed that ezrin enrichment in the brush border was lost within a 3-hour window following exposure to NSC668394. Notably, in all these events, the loss of ezrin signal was followed by a striking increase of MISP and UtrCH signal throughout the brush border (**Fig. 6D, E; Movie S3**). Moreover, overaccumulation of MISP and UtrCH in NSC668394-treated W4 cells also coincided with a dramatic increase in microvillar length (**Fig. 6D; UtrCH channel**). However, these instances of elongation were temporary as protrusions eventually collapsed after 30-60 min of growth without impacting the accumulation of MISP and UtrCH at the base of the brush border. To further define the impact of ezrin accumulation on MISP localization and microvillar structure, we used SIM to take a closer look at the brush border of W4 cells fixed after 2 hours of NSC668394 treatment. SIM images revealed a significant increase in the overall length of core actin bundles in NSC668394-treated cells compared to control cells (3.58 ± 0.83 μm vs. 8.25 ± 2.11 μm) (**Fig. 6F, G**). Interestingly, MISP occupancy along core bundles also increased from 38% in DMSO-treated cells to 53% in NSC668394-treated cells (**Fig. 6H**). These findings indicate that ezrin and its associated membrane-actin linking activity confine MISP to the rootlets of microvilli.

## DISCUSSION

The filament crosslinking activity of actin bundling proteins provides the microvillar core bundle with the flexural rigidity needed to overcome plasma membrane tension and protrude from the cell surface (Atilgan et al., 2006). Although villin, fimbrin, and espin are canonical bundling proteins that have been identified and characterized in the context of the epithelial brush border, the persistence of microvillar growth in mice lacking all three of these factors suggested the existence of as-of-yet unidentified bundlers (Revenu et al., 2012). Epidermal growth factor receptor pathway substrate 8 (EPS8) has been invoked as a fourth bundler with the potential to compensate for crosslinking activity in the absence of other canonical crosslinkers (Revenu et al., 2012). However, its exquisitely specific localization to the distal tips of microvilli is at odds with the need for canonical crosslinkers to be distributed along the length of the core bundle. Additionally, whereas certain cell types employ isoforms of fascin to drive robust parallel bundling of filaments in other related actin-based protrusions such as filopodia and stereocilia (Krey et al., 2016; Roy and Perrin, 2018; Svitkina et al., 2003), there is no evidence for fascin expression in transporting epithelia of the gut and kidney. Thus, the identity of other functional bundlers in the apical brush border has remained an open question.

Here, we identify MISP as a novel component of the epithelial brush border that holds actin filament bundling potential. Previous studies on MISP have focused on its role during mitotic progression in nonepithelial cells (Maier et al., 2013; Zhu et al., 2013). In that context, MISP localizes to the actin rich cortex and contributes to anchoring the spindle by interacting with microtubule asters during metaphase (Maier et al., 2013; Zhu et al., 2013). In native intestinal tissues and differentiating epithelial cell culture models, we found that MISP localizes to the base of core actin bundles that support microvilli. Moreover, careful examination of confocal and super-resolution images revealed that MISP is restricted to the rootlet ends of core bundles, which are embedded in the subapical terminal web and thus are not wrapped by plasma membrane. Such a pattern of localization is consistent with previous studies showing MISP enrichment in the proximal region of the neuronal growth cone, where the pointed ends of filopodial actin filaments coalesce (Kumeta et al., 2014). Thus, localization near pointed ends of actin filaments appears to be a conserved property of MISP. Interestingly, MISP labeling is observed not only in the terminal web of enterocytes along the villus, but also in the subapical region of immature/differentiating enterocytes found in the crypt. Therefore, MISP is enriched at the cell apex during the window of differentiation when microvilli are actively growing. Having a rootlet specific bundler present at early stages of differentiation is consistent with classic ultrastructural studies, which suggested that the growth of new microvilli is supported by a simultaneous maturation of the terminal web immediately beneath the apical membrane (Tilney and Cardell, 1970).

Our data indicate that MISP functions in the selective stabilization of the rootlet ends of core bundles as KD in W4 cells results in significant shortening of rootlets when visualized with super-resolution microscopy. In contrast, overexpression of MISP promoted a moderate but significant elongation of rootlets. These phenotypes are explained by MISP’s filament bundling activity, which we reconstituted *in vitro*. Actin filament bundles assembled with purified MISP demonstrate tight packing with an average inter-filament spacing of ~10.2 nm, which is close to that reported for fimbrin (~9-12 nm) (Matsudaira et al., 1983; Volkmann et al., 2001), but slightly shorter to the distance between filaments bundled by villin or espin (~12 nm) (Bartles et al., 1998; Hampton et al., 2008). This suggests that the arrangement of filaments in intact microvilli reflects the collective activity of multiple bundling proteins, each bringing their own characteristic spacing. Indeed, previous EM studies on filament packing and spacing in stereocilia core bundles, which are occupied by fascin-2, espin-1, and fimbrin (Krey et al., 2016) and TRIOBP-4 and −5 (Kitajiri et al., 2010) are consistent with this general idea.

Intriguingly, co-expression of MISP and fimbrin cooperatively elongated core bundle rootlets deep into the cytoplasm of W4 cells. The exaggerated nature of these rootlets allowed us to capture the temporal details of their formation, which preceded the assembly of microvillar protrusions. Thus, the apical localization of bundlers early in enterocyte differentiation might provide mechanical stability to nascent and growing microvilli. Ectopic expression experiments in HeLa cells that generally do not make microvilli also revealed that MISP can drive the formation of aberrant actin bundles and these structures, in turn, recruit fimbrin. Yet how MISP binds to and bundles actin filaments and recruits fimbrin remains unknown. In our analysis, we were unable to identify recognizable actin binding and bundling motifs in the MISP primary sequence, although previous studies point to multiple actin binding motifs distributed throughout the molecule (Kumeta et al., 2014), which is consistent with functional requirements of a bundling protein. Considering the cooperative effects of MISP and fimbrin on rootlet length and stability, it is tempting to speculate that these factors bind to different sites on actin filaments. In contrast to MISP, the multiple actin binding domains (ABDs) of fimbrin are well characterized (Klein et al., 2004), and their binding sites on F-actin in 2D arrays have been mapped using cryo-EM (Volkmann et al., 2001). Based on these structural studies, we speculate that MISP binds outside the canonical inter-monomer cleft that is targeted not only by fimbrin but also cofilin (Tanaka et al., 2018), myosin (Mentes et al., 2018), and even live imaging probes such as Lifeact (Belyy et al., 2020). Future cryo-EM studies aimed toward elucidating the structural details of the MISP binding site on F-actin will be needed before we can begin to understand the nature of MISP/fimbrin cooperativity. Independent of a detailed actin binding and bundling mechanism, the hierarchical targeting of MISP and fimbrin suggests an order of action for these two bundling proteins during microvillar assembly. We propose that MISP expression and localization to rootlets leads the arrival of fimbrin at the apical surface in differentiating enterocytes. To examine this possibility during microvilli biogenesis, high temporal resolution live imaging studies with differentiating epithelial cells using recently developed approaches will be needed (Gaeta et al., 2021).

The highly restricted targeting of MISP to the rootlet is unique among epithelial actin bundlers, though previous studies revealed that fimbrin accumulates at higher levels at the proximal ends of core bundles in the terminal web, relative to the distal end (Grimm-Günter et al., 2009). In MISP KD cells, we noted that the segment of the core actin bundle enveloped in plasma membrane increased in parallel with the shortening of rootlets induced by loss of MISP. This finding suggested a previously unrecognized interplay between mechanisms that control the extent of membrane wrapping (i.e. the length of protruding microvillus) and the activity of actin bundling proteins that dictate rootlet length. Thus, we hypothesized that factors that simultaneously bind to plasma membrane and actin would be well positioned to prevent MISP binding along the more distal membrane-wrapped segment the core actin bundle. A common feature of actin-based protrusions is the tethering of cytoskeleton to the enveloping membrane by ERM (ezrin, radixin, moesin) proteins (Revenu et al., 2004), with ezrin being the most abundant ERM in the brush border. Remarkably, inactivation of ezrin using a small molecule inhibitor led to the release of ezrin from the plasma membrane and immediate ectopic redistribution of MISP from the rootlets up to more distal regions of the core bundle. Loss of active ezrin and redistribution of MISP also led to a drastic increase in microvillar length. This striking response might reflect loss of the mechanical constraint that the membrane normally imposes on the distal barbed ends, the preferred site of actin monomer incorporation. Alternatively, MISP recruitment to more distal regions of the bundle might directly promote stabilization and slowing of the robust treadmilling and turnover that normally occur in this system (Meenderink et al., 2019; Tyska and Mooseker, 2002). Notably, while high ezrin levels drastically displaced MISP from rootlets, increasing MISP levels had no impact on the membrane-wrapped segment of core actin bundles. This further suggests that ezrin exerts a dominant effect on MISP confinement to core bundle rootlets. Taken together, these data argue for a mutual exclusivity model where opposite ends of core actin bundles are decorated by either ezrin or MISP and the balance between these populations ultimately dictates the extent of membrane coverage.

The fact that ezrin excludes MISP from binding along the membrane-wrapped segment of the microvillus may also offer additional insight on where MISP resides in a core bundle. If one assumes that membrane-associated ezrin only binds to actin filaments superficially exposed on the surface of the core bundle, MISP’s inability to occupy distal regions might suggest that this bundler also binds superficially. Although speculative, such superficial binding has been demonstrated for TRIOBP-4, a bundling protein that targets specifically to the rootlets of hair cell stereocilia (Kitajiri et al Cell 2010). MISP and TRIOBP do not share motifs or domain organization, but secondary structure analysis using Phyre2 predicted that MISP sequence is largely disordered as it has been reported for TRIOBP-4 (Bao et al., 2013). Thus, it remains possible that MISP bundles filaments using a similar mechanism. It is also worth noting that other well characterized actin bundlers in microvilli – villin and espin – are not restricted to the surface of core bundles and also do not exhibit mutually exclusive localization with ezrin.

In conclusion, the discoveries reported here point to a new mechanism for bundling actin filaments, and specifically those that comprise the core bundle rootlets of epithelial microvilli. These findings strengthen our molecular understanding of the biologically robust formation of evolutionary conserved microvillus-rich apical specializations. Indeed, the emergence of MISP as a new linear actin bundler may offer a molecular explanation for the remarkable finding that triple villin-espin-fimbrin KO mice are still capable of assembling brush border microvilli (Revenu et al., 2012). As MISP is also implicated in promoting mitotic progression (Maier et al., 2013; Zhu et al., 2013), our discoveries further suggest a potential role of MISP in coupling oriented cell division with differentiation in transporting epithelial cells.

## MATERIAL AND METHODS

### Cell culture

Ls174T-W4 cells (W4; human colon epithelial cancer cells), CACO-2_BBE_ cells (human colorectal adenocarcinoma epithelial cell line), LLC-PK1-Cl4 cells (CL4; pig kidney epithelial cells), HeLa cells (human cervical cancer cell line), and HEK293T cells were cultured in Dulbecco’s modified Eagle’s (DMEM) medium with high glucose and 2 mM L-glutamine (Corning; 25-005-CI). Ls174T-W4 cells (a generous gift from Dr. Hans Clevers) were grown in media supplemented with 10% tetracyclin-free fetal bovine serum (Atlanta Biological, S10350), 1 mg/ml G418 (Gold Biotechnology; G-418), 10 *μ*g/ml blasticidin (Gold Biotechnology; B-800), and 20 *μ*g/ml phleomycin (InvivoGen; ant-ph-1). For CACO-2_BBE_ cells, the media was supplemented with 20% fetal bovine serum. For LLC-PK1-Cl4, HeLa, and HEK293T cells, the media was supplemented with 10% fetal bovine serum. All cultured cells were grown at 37 °C and 5% CO_2_.

### Cloning and constructs

The human MISP sequence harbored in a pCMV-SPORT plasmid (Harvard PlasmID Database; HsCD00326629) was subcloned by PCR and TOPOcloned into a pCR™8 Gateway entry vector (Invitrogen; 46-0899). In-frame sequence insertion was confirmed by sequencing. MISP was then shuttled into Gateway-adapted plasmids: pEGFP-C1, pmCherry-C1, and pHALO-C1. Similarly, the human beta-actin and UtrCH sequences were cloned and shuttled into a Gateway adapted HALO-C1 plasmid. The human fimbrin sequence was cloned into a pEGFP-C1 plasmid (Clontech; 6084-1). The human ezrin sequence was cloned into a pEGFP-N1 (Clontech; 6085-1). The human MISP and EGFP-MISP sequences were subcloned into modified pFastBac-6xHis-MBP plasmid LIC expression vector (Addgene; plasmid #30116). All constructs were confirmed by sequencing.

### Transfection and lentivirus production

For overexpression experiments, cells were transfected using Lipofectamine2000 (Invitrogen; 11668019) according to the manufacturer’s instructions. For KD experiments, lentiviral particles were generated by transfecting HEK293T cells with 6 *μ*g PLKO.1 scramble control and MISP-targeted shRNA plasmids (SIGMA; TRCN0000422523, TRCN0000116527), 4 *μ*g psPAX2 packing plasmid (Addgene, 12260), and 0.8 *μ*g pMD2.G envelope plasmid (Addgene; #12259) using FuGENE 6 (Promega; E2691). Lentiviral particles were harvested and concentrated using a Lenti-X Concentrator (Clontech; 631231). Concentrated lentiviral particles were supplemented with polybrene (SIGMA; H9268) and incubated with W4 cells at 80% confluency. After 24 hours, the media was replaced with fresh media containing puromycin (Gold Biotechnology; P-600-100) for selection. Selection was applied for 14 days, replacing with fresh selection media every other day. Rescue experiments were conducted using an EGFP-MISP construct designed to be refractory to shRNA KD.

### Immunofluorescence

Cells grown on a glass coverlips were fixed with 4% paraformaldehyde (EMS; 15710) in 1X PBS for 15 minutes at 37 °C. Fixed cells were washed with 1X PBS, and permeabilized with 0.1% Triton X-100 in 1X PBS for 15 minutes at room temperature. Cells were washed with 1X PBS and blocked with 5% Bovine Serum Albumin (BSA) in 1X PBS for 2 hours at room temperature. Cells were washed and incubated with primary antibodies overnight at 4 °C. Primary antibodies used were anti-MISP (Thermo Scientific; PA5-61995), anti-villin (Santa Cruz; sc-66022), anti-ezrin (CST; 3145), anti-EPS8 (SIGMA; HPA003897). Cells were washed with 1X PBS four times for 5 minutes and incubate with secondary antibodies. Goat anti-rabbit Alexa Fluor 488 F(ab’)2 Fragment (Molecular Probes; A11070), goat anti-mouse Alexa Fluor 568 F(ab’)2 Fragment (Molecular Probes; A11070), Alexa Fluor 568-phalloidin (Invitrogen; A12380), Wheat Germ Agglutinin 405M (WGA) (Biotium; 29028-1), DRAQ5 (Thermo Scientific; 62251). Cells were washed again with 1X PBS and mounted on glass slides using ProLong Gold (Invitrogen; P36930).

### Western blot analysis

Cell lysates were prepared using RIPA buffer (SIGMA; R0278) supplemented with protease inhibitors (Roche; 04693124001). Samples were centrifuged at 20,000 × *g* for 15 minutes to remove cell debris. The resulting supernatant was boiled with Laemmli sample buffer for 5 minutes. Samples were then loaded on a 4-12% NuPAGE gradient gel (Invitrogen; NP0322BOX). Gels were transferred onto a nitrocellulose membrane at 30V for 18 hours. Membranes were blocked with 5% dry milk diluted in 1X PBS containing 0.1% Tween-20 (PBS-T) for 2 hours at room temperature. The membranes were incubated with primary antibody diluted in 1X PBS-T containing 1% BSA overnight at 4C. Primary antibodies used were anti-MISP (Thermo Scientific; PA5-61995), anti-villin (Santa Cruz; sc-66022), anti-GAPDH (Cell Signaling; 2118), anti-β-actin (SIGMA; A5316). Membranes were then washed with 1X PBS-T and incubated with secondary antibodies for 1 hour at room temperature. Secondary antibodies used were IRdye 800 donkey anti-rabbit (LI-COR; 926-32213) or donkey anti-mouse (LI-COR; 926-32212). Membranes were washed with 1X PBS-T and imaged using the Odyssey CLx infrared scanner (LI-COR). Images were processed using the FIJI software (NIH). Protein expression levels were normalized to GAPDH.

### Light Microscopy and image processing

Laser scanning confocal imaging was conducted using Nikon A1 Microscope equipped with 405, 488, 561 and 645 nm LASERs, Apo TIRF 100x/1.45 NA, Plan Apo 60x/1.4 NA, Plan Fluor ELWD 40x/0.6 NA objectives. Live-cell imaging was conducted using a Nikon Ti2 Eclipse equipped with 488, 561 and 645 nm excitation LASERs, Apo TIRF 100x/1.49 NA and Plan Fluor 40x/1.3 NA objectives, a Hamamatsu X1 spinning disk, and Photometrics Prime 95B sCMOS or Hamamatsu Orca-Fusion BT sCMOS cameras. FRAP was also conducted using a Bruker mini-scanner module capable of producing ROI specific 405 nm photo-stimulation. Images were deconvolved and/or denoised using Nikon Elements software. Super-resolution imaging was performed using a Nikon Structured Illumination Microscope (N-SIM) equipped with 405, 488, 561 and 640 nm LASERs, an SR Apo TIRF 100x/1.49 NA objective, and an Andor iXon Ultra DU-897 EMCCD camera. Images were reconstructed using Nikon Elements software. For imaging in all microscope modalities, gain was matched between samples during image acquisition.

### Protein purification

6xHis-MBP-MISP and 6xHis-EGFP-MBP-MISP constructs were expressed in Sf9 insect cells. Insect cell pellets were resuspended in lysis buffer (20 mM Tris HCl, 0.3 M KCl, 10 mM imidazole, 10% glycerol, 2 mM DTT, pH 7.5) supplemented with protease inhibitors (Roche, 5892953001). Resuspended samples were lysed using a Dounce homogenizer and passed through an 18-gauge needle to shear DNA. The resultant lysate was then centrifuged at 35,000 rpm in a Ti 50.2 rotor (Beckman) for 30 minutes at 4 °C. Clarified lysates were then filtered using a 0.45 *μ*m syringe filter. Samples were then loaded into a HisTrap column according to the manufacturer protocol and eluted with a 50-500 mM linear imidazole gradient (pH 7.5). Protein purity was assessed by SDS-PAGE. Eluted protein was concentrated using a centrifugal filter (Millipore; UFC803024). For *in vitro* EM experiments, the 6xHis-MBP tag was cleaved from 6xHis-MBP-MISP using a TEV protease (NEB; P8112) for 1 hour at room temperature. The cleaved 6xHis-MBP tag was removed by incubating the solution with Ni-NTA magnetic beads (NEB; S1423) for 1 hour at 4 °C. The solution was then placed in a magnetic rack to separate the bead-bound 6xHis-MBP fraction from MISP. The purified full-length MISP was run in an SDS-PAGE gel to confirm successful cleavage.

### Low Speed Sedimentation Assay

Rabbit skeletal G-actin (Cytoskeleton Inc., AKL99) was resuspended according to manufacturer instructions. Resuspended G-actin were centrifuged at 100,000 × *g* to remove aggregated monomers. G-actin was then polymerized according to the manufacturer instruction. F-actin was stabilized with phalloidin (SIGMA, A22287), and centrifuged at 20,000 × *g* to precipitate nonspecific aggregates. F-actin (5 *μ*M) was incubated with increasing concentrations of 6xHis-MBP-MISP (0–5 *μ*M) for 15 minutes at room temperature. Subsequently, all MISP/F-actin sample series were centrifuged at 10,000 × *g* for 20 minutes at room temperature. The supernatant was carefully removed without disrupting the pellet. All pellets were boiled with samples buffer and run into a 4-12% NuPAGE gradient gel (Invitrogen, NP0322BOX). Gels were stained with Coomassie blue (Bio-Rad, 1610786) and imaged in a gel imaging system (Bio-Rad, Gel Doc™ EZ System).

### Transmission Electron Microscopy

To prepare MISP/F-actin mixtures for electron microscopy (EM), F-actin was prepared as previously described. Phalloidin-stabilized F-actin was incubated with or without purified MISP at a 5:1 molar ratio overnight at 4 °C. Carbon-coated copper grids (EMS; cat# CF300-Cu) were glow discharged and coated with 0.1% poly-lysine solution for 15 min and washed 2 times with ddH_2_O to remove free poly-lysine. Samples were incubated with the grids for 15 min, briefly washed with ddH_2_O, and negative stained with 2% uranyl acetate. Images were collected on a FEI Tecnai T-12 transmission electron microscope operating at 100 kV using an AMT CMOS camera.

### Image analysis and statistical testing

All images were process and analyzed using Nikon Elements software or FIJI software package (https://fiji.sc). Time series volumes from live imaging experiments were registered using the StackReg plugin in FIJI as needed.

#### Analysis of signal intensity in intestinal tissue samples

To measure signal intensities along microvilli, a 1-pixel-wide line scans were drawn along the base-tip axis of brush border. To measure signal intensities in the brush border along the crypt-villus axis, an ROI containing the brush border was drawn and straightened using the Straighten plugin in FIJI; averaged intensities were calculated across the resulting rectangle. All intensity values were normalized from 0 (base) to 1 (tip) and fit to a Gaussian curve using PRISM v. 9.0.

#### Analysis of brush border assembly in W4 cells

To quantify the percentage of W4 cells capable of forming brush borders, cells exhibiting a single polarized cap of F-actin (representing a brush border) were counted manually. For rescue experiments, only W4 cells expressing an EGFP-MISP refractory construct were scored. To quantify the overall actin intensity in W4 cells, multiple ROIs containing single cells were generated using Nikon Elements software, and F-actin intensities measured in each ROI.

#### Measuring the lengths of microvilli and rootlets in W4 cells

For the purposes of quantification throughout the paper, we define a microvillus as the segment of a core bundle that is wrapped in plasma membrane, and ‘rootlet’ as the segment that is free of membrane wrapping. Microvilli and rootlet lengths were measured separately using a membrane marker to delineate the boundary of these regions. To calculate membrane coverage (i.e. fraction of the core bundle wrapped in membrane), we summed the averaged lengths of microvilli and rootlets per cell to obtain a total core bundle length. We then calculated membrane coverage as the ratio of average microvilli length to total core bundle length.

#### Inter-filament spacing

To quantify the spacing between MISP-bundled actin filaments, EM images were process using the FFT bandpass filter in FIJI. We used image filtering to remove small structures down to 10 pixels (5 nm) and large structures up to 100 pixels (50 nm). Line scans were then drawn perpendicular to tightly packed actin filaments, signal intensity was plotted, and the distance between peaks was measured. For control conditions, actin bundles with an inter-filament spacing of less than 20 nm were considered for quantification and processed as described above.

#### FRAP analysis

ROIs of similar area were drawn over the microvilli and rootlets of W4 cells, and bleached using a 405 nm LASER steered with a Bruker mini-scanner. Cells were imaged for 30 seconds before photobleaching, bleached over the course of 5 sec, and then imaged every 10 sec for 30 min to capture signal recovery dynamics. All intensity values for each condition were normalized from 0 to 1 and plotted together to facilitate comparison. Averaged values for each condition were fit using two-phase association curves.

#### Microvilli and rootlet assembly in W4 cells

To quantify the intensity of actin turnover in the microvilli and rootlets in W4 cells overexpressing mCherry-MISP, EGFP-fimbrin and HALO-β-actin, we drew ROIs delimiting these domains in the β-actin channel. As cells were not synchronized in their differentiation following doxycycline addition, in each cell we set ‘0’ as the time frame where the β-actin signal in the rootlet domain increased above background.

#### Ezrin inhibition in W4 cells

To quantify MISP enrichment to the brush border upon ezrin inhibition in W4 cells overexpressing mCherry-MISP, EGFP-ezrin, and HALO-UtrCH, we drew ROI containing the brush border in the β-actin channel. As the effect of NSC668394 on ezrin inhibition from the brush border was not synchronized across cells, we arbitrarily set the time point ‘0’ as 9 frames (22.5 minutes) before ezrin signal dropped below background. All intensity values for each condition were normalized from 0 to 1 and plotted together to facilitate comparison.

### Statistical analysis

Statistical significance was performed using the unpaired T-test for pairwise comparison. Statistical correlation was conducted using the Pearson correlation coefficient for colocalization analysis. All statistical analysis was computed in PRISM v. 9.0. (GraphPad).

## ACKNOWLEDGEMENTS

The authors would like to thank all members of the Tyska laboratory, members of the Vanderbilt Microtubules and Motors Club, and the Epithelial Biology Center for feedback and advice. Microscopy was performed in part by the Vanderbilt Cell Imaging Shared Resource. This work was in part supported by NIH grants R01-DK111949, R01-DK095811, and R01-DK125546 (MJT). MZ acknowledges the support by NIH grant R35-GM119552.

## AUTHOR CONTRIBUTIONS

EAM and MJT designed the experiments. EAM performed experiments, analyzed the data, and prepared the figures. CA, MZ, ESK assisted with provided expertise with experimental methodology, and contributed significant conceptual insight. EAM and MJT wrote the manuscript. All authors contributed to revising the manuscript.

## SUPPLEMENTARY INFORMATION

**Figure S1.**
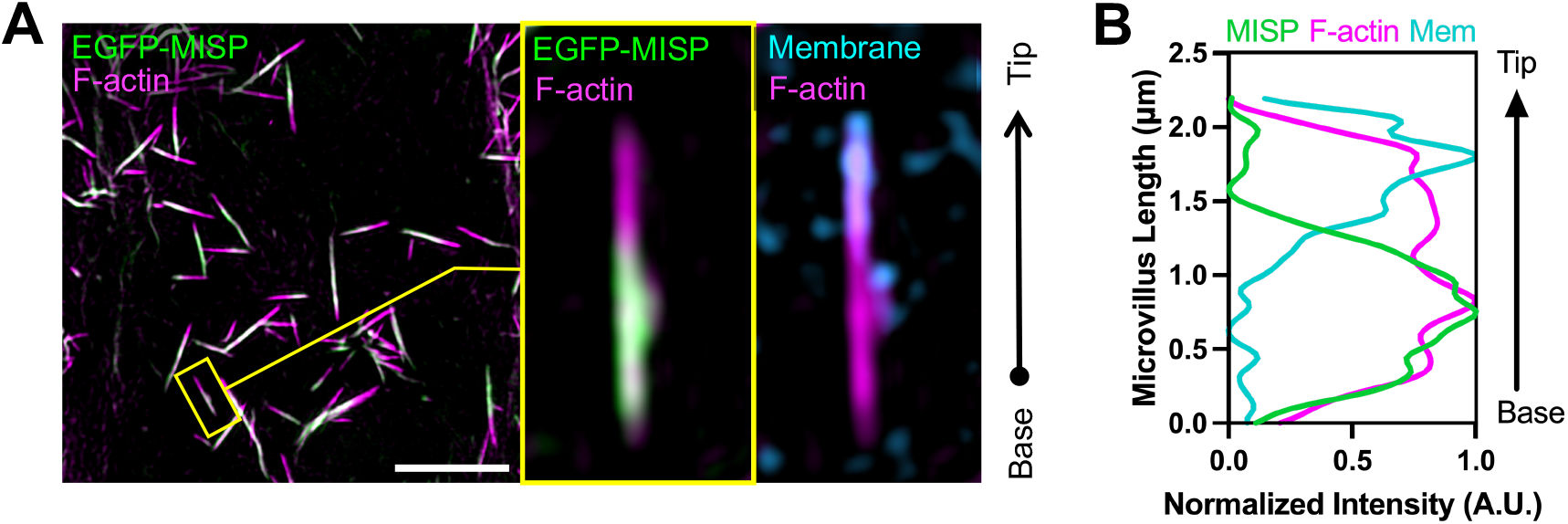
related to Figure 1. **(A)** SIM maximum intensity projection image of a CL4 overexpressing EGFP-MISP and stained for F-actin with phalloidin (magenta); and membrane with WGA (cyan). The left panel shows combined channels. The right panel shows zoomed images of the yellow box shown in the left panel. Scale bar = 5 *μ*m. **(B)** Fluorescence intensity measurements of an individual microvillus shown in the zoomed image in D.

**Figure S2.**
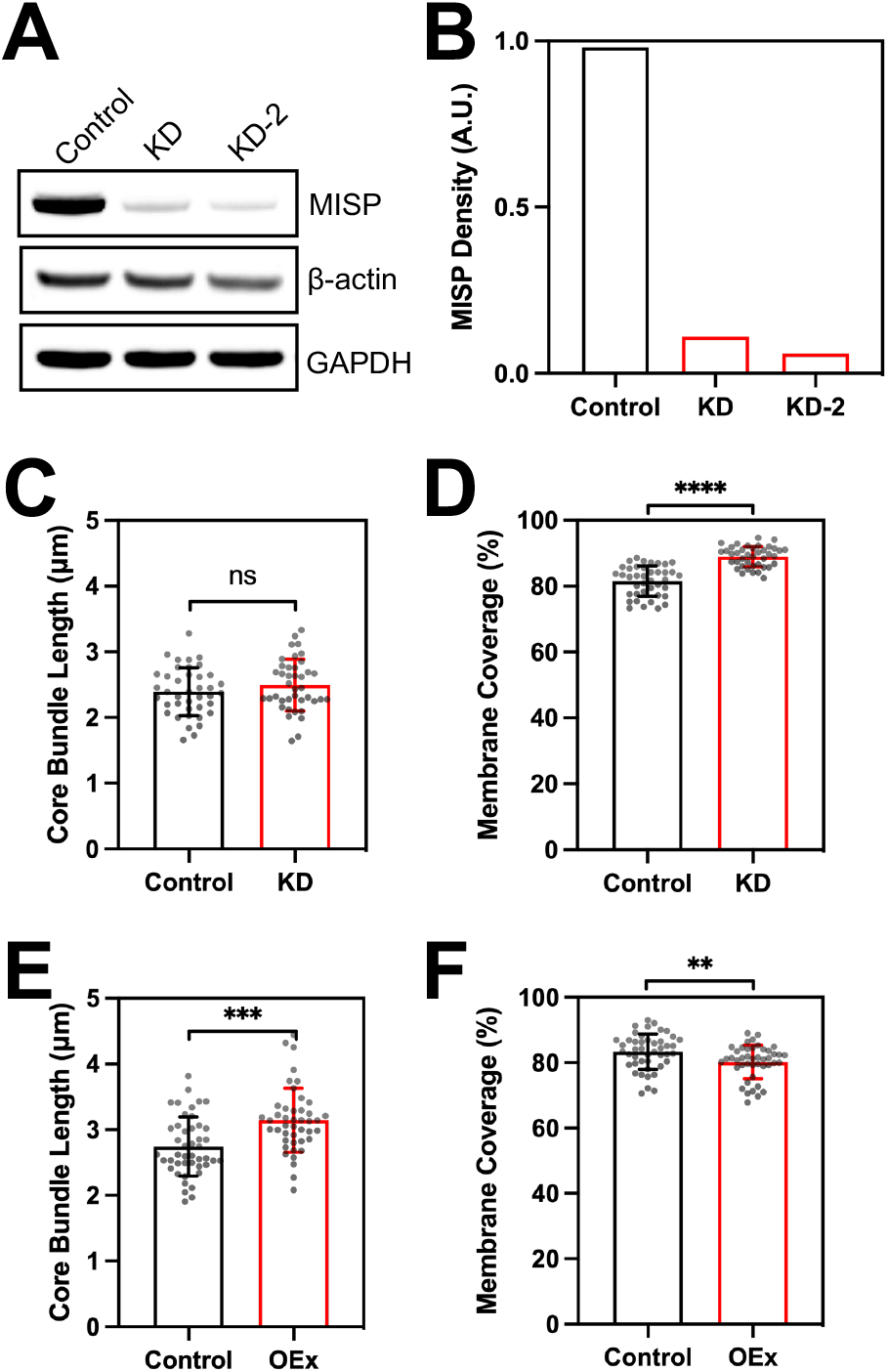
related to Figure 2. **(A)** Western blot analysis of endogenous MISP in scramble control and MISP KD W4 cells. **(B)** Density quantification of MISP bands from western blot shown in A. Densities were normalized to GAPDH. **(C, E)** Length measurements of core actin bundles for KD (C) and overexpression (E) conditions described in Figure 2D and 2F, respectively. Each data point represents the average length of > 10 core actin bundles per cell; n ≥ 40 cells per condition. All data points are representative of three independent experiments. **(D, F)** Membrane coverage measurements of core actin bundles for KD (D) and overexpression (F) conditions described in Figure 2D and 2F, respectively. Each data point represents the membrane coverage percentage of core actin bundles per cell; n ≥ 40 cells per condition. All data points are representative of three independent experiments. All bar plots and error bars denote mean ± SD. p-values were calculated using the unpaired T-test (ns: not significant; **: p < 0.01; ***: p < 0.001; ****: p < 0.0001).

**Figure S3.**
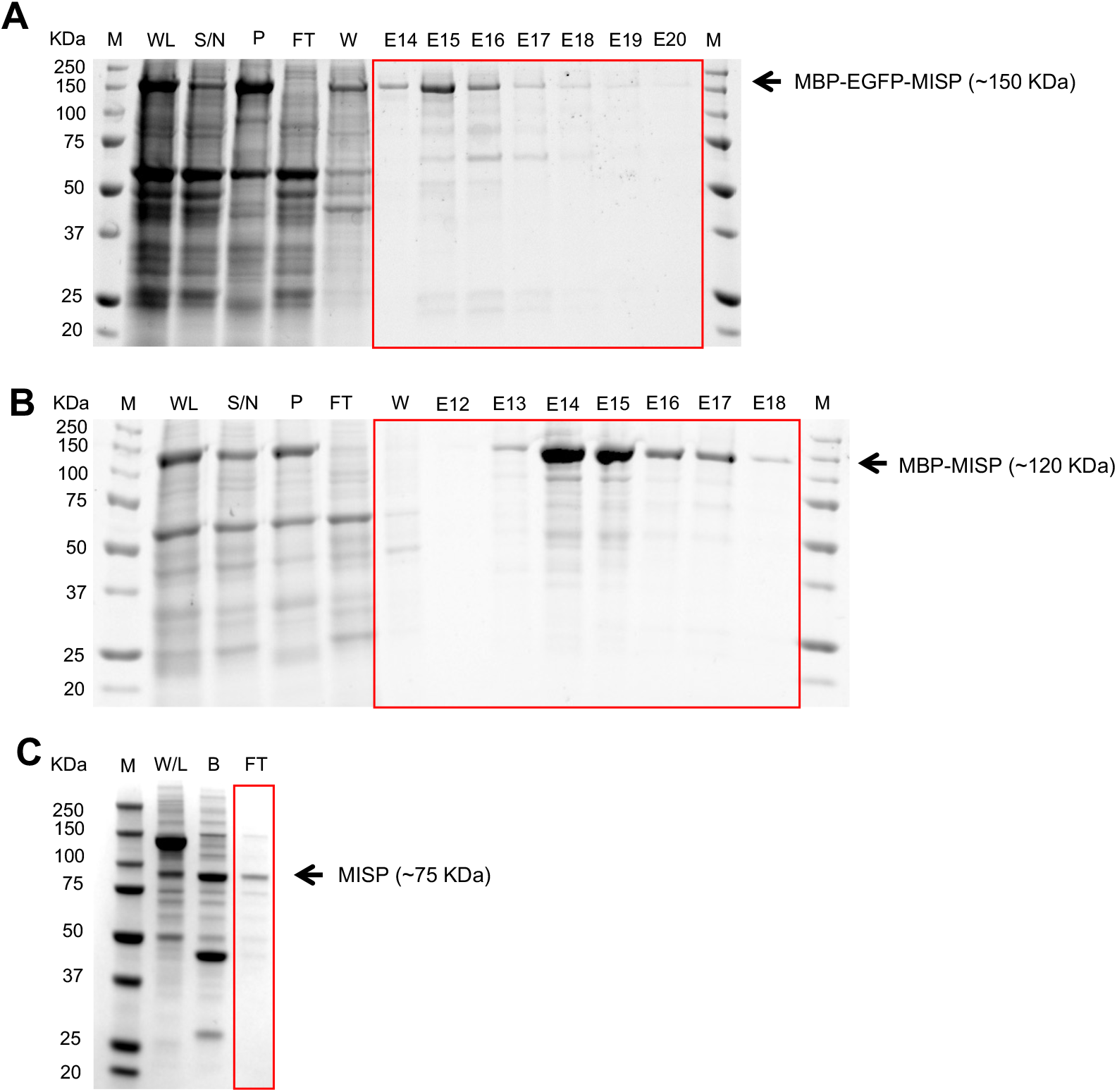
related to Figure 3. **(A-B)** Purification of MBP-EGFP-MISP and MBP-MISP from Sf9 insect cells. Coomassie-stained SDS polyacrylamide gel showing all purification steps: whole lysate (WL), supernatant fraction (S / N), pellet fraction (P), flow through (FT), wash (W), and elution (E). ‘M’ denotes protein ladder marker (10-250 KDa). **(C)** TEV protease cleavage of purified MBP-MISP. Coomassie-stained SDS polyacrylamide gel showing purification steps: control whole lysate (NC), fraction bound to beads after TEV cleavage (B), flow through (FT). Purified MISP was recovered from the flow through fraction. ‘M’ denotes protein ladder marker (10-250 KDa).

**Figure S6.**
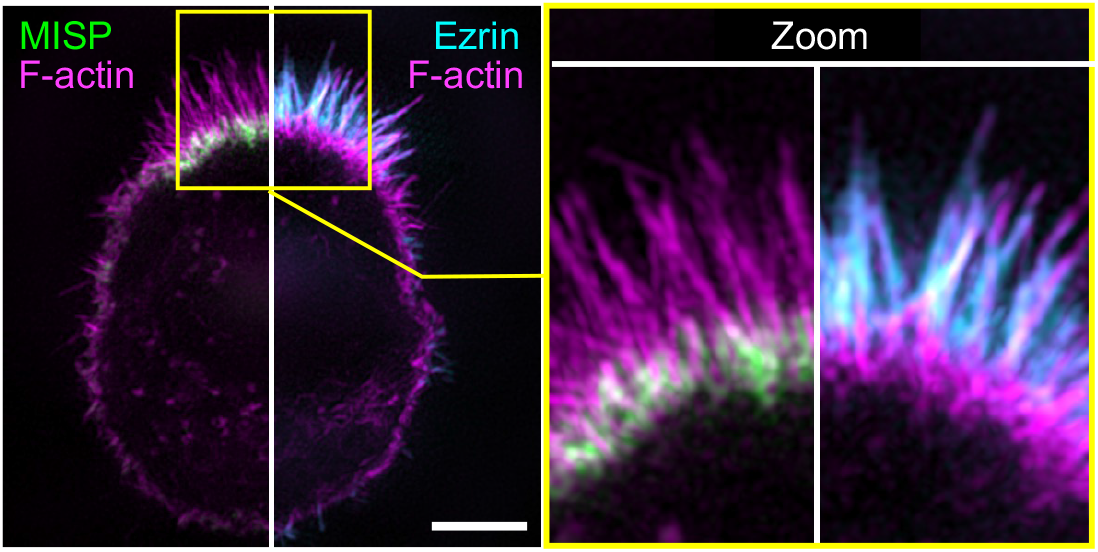
related to Figure 6. **SIM** maximum intensity projection image of a W4 cell stained for: MISP (green); ezrin (cyan); and F-actin with phalloidin (magenta). The left panel shows the image split into two to display a combination of two-color channels. The right panel shows zoomed images of the yellow box shown in the left panel. Scale bar = 3 *μ*m.

**Movie S1, related to Figure 5H.** The assembly of hyper-elongated rootlets precedes the formation of protruding microvilli. Confocal maximum intensity projection movie of W4 cells overexpressing HALO-β-actin, EGFP-fimbrin and mCherry-MISP after the addition of doxycycline. Inverted single channels are shown in the left panels, and the merged channel is shown in the right panel. Scale bar = 10 *μ*m

**Movie S2, related to Figure 6A.** Ezrin overexpression displaces endogenous MISP to the brush border periphery creating a ring-like distribution of MISP around ezrin. SIM maximum intensity projection movie of a fixed W4 cell overexpressing EGFP alone (left cell) or EGFP-ezrin (right cell). Cells were also stained for endogenous MISP (magenta). Scale bar = 5 *μ*m

**Movie S3, related to Figure 6D.** Ezrin inhibition by NSC668394 promotes MISP accumulation to more distal segments of core actin bundles. Spinning disk confocal maximum intensity projection movie of a W4 cell overexpressing EGFP-ezrin (green), mCherry-MISP (magenta), HALO-UtrCH (cyan). Inverted single channels are shown in the left panels, and the merged channel is shown in the right panel. Scale bar = 5 *μ*m

